# LRP1 is an entry receptor for the botulinum toxin complex in the gut

**DOI:** 10.64898/2026.06.25.734645

**Authors:** Sho Amatsu, Takuhiro Matsumura, Chiyono Morimoto, Hideo Yagita, Kiyo-aki Ishii, Takashi Kanaya, Koji Hase, Nobuhide Kobayashi, Masahiko Zuka, Hiroshi Ohno, Toshinari Takamura, Yukako Fujinaga

**Affiliations:** Department of Bacteriology, Graduate School of Medical Sciences, Kanazawa University, Ishikawa 920-8640, Japan; Department of Forensic Medicine and Pathology, Graduate School of Medical Sciences, Kanazawa University, Ishikawa 920-8640, Japan; Department of Immunology, Juntendo University Graduate School of Medicine, Tokyo 113-8421, Japan; Department of Endocrinology and Metabolism, Kanazawa University, Ishikawa 920-8640, Japan; Laboratory for Intestinal Ecosystem, RIKEN Center for Integrative Medical Sciences, Kanagawa 230-0045, Japan; Graduate School of Medical Life Science, Yokohama City University, Kanagawa 230-0045, Japan; Division of Commensal Biology, The Institute of Medical Science, The University of Tokyo, Tokyo 108-8639, Japan; International Vaccine Design Center, The Institute of Medical Science, The University of Tokyo, Tokyo 108-8639, Japan; Division of Biochemistry, Faculty of Pharmacy and Graduate School of Pharmaceutical Science, Keio University, Tokyo 105-8512, Japan; Institute of Fermentation Sciences (IFeS), Faculty of Food and Agricultural Sciences, Fukushima University, Fukushima 960-1296, Japan

**Keywords:** LRP1, low-density lipoprotein receptor–related protein 1, *Clostridium botulinum*, botulinum toxin complex, hemagglutinin, high-oral-toxic, transcytosis, uptake, enterocyte

## Abstract

Botulinum neurotoxin (BoNT) is an etiologic agent of food poisoning caused by *Clostridium botulinum*. The large progenitor toxin complex (L-PTC) crosses the intestinal epithelial barrier to deliver BoNT to target neurons; however, it is not clearly understood how BoNT enters the host. Here, we identified low-density lipoprotein receptor–related protein 1 (LRP1) as a major enterocyte transcytosis receptor for the hyper-oral-toxic L-PTC serotype B-Okra (L-PTC/B^Okra^). We found that hemagglutinin (HA), a neurotoxin-associated protein within the L-PTC/B^Okra^ complex, binds to LRP1 via *N*-glycans. HA/B^Okra^ co-localized with LRP1 within the internalized vesicles in cultured cells and enterocytes. LRP1 deletion inhibited the apical-to-basal transcytosis of L-PTC/B^Okra^ in an intestinal epithelial cell line, and this effect was rescued by LRP1 re-expression. Finally, intestinal epithelial cell–specific LRP1-deficient mice displayed reduced susceptibility to toxicity caused by oral administration of L-PTC/B^Okra^. Taken together, these results indicate that *N*-glycosylated LRP1 mediates L-PTC/B^Okra^ transcytosis via enterocytes, enabling BoNT to traverse the intestinal epithelial barrier.

## Introduction

Botulism is a rare but life-threatening disease that causes neuroparalysis due to a neurotoxin (BoNT) produced by *Clostridium botulinum* and related clostridial species^1^. BoNT cleaves soluble *N*-ethylmaleimide-sensitive-factor attachment protein receptor (SNARE) proteins that are required to release acetylcholine into neuromuscular junctions, resulting in flaccid paralysis. Foodborne botulism, a common form of the disease, results from oral BoNT intake. After the ingestion of contaminated food, BoNT is absorbed through the gastrointestinal tract into the bloodstream and peripheral nerve terminals^2^.

BoNT is naturally produced as a component of progenitor toxin complexes (PTCs) and neurotoxin-associated proteins (NAPs), the latter of which include non-toxic non-hemagglutinin (NTNHA) and hemagglutinin (HA)^2^. Some BoNT-producing strains carry the *orfX* gene cluster instead of the *ha* gene cluster^3^, although their molecular functions remain unclear. BoNT and NTNHA form medium PTC (M-PTC), which protects BoNT from gastric degradation^4,5^. M-PTC associates with HA to form large PTC (L-PTC), which facilitates BoNT absorption in the small intestine^6,7^. HA increases the toxicity of orally administered M-PTC by 30-to 700-fold relative to M-PTC alone^2^. BoNTs have traditionally been classified into seven serotypes (A–G); serotypes A, B, and E cause human botulism, whereas F rarely does^2^. In mice, L-PTC serotype B-Okra (L-PTC/B^Okra^) exhibits at least 20- to 80-fold higher toxicity after oral administration than serotype A-62A (L-PTC/A^62A^)^2,8^. We recently proposed that L-PTCs can be divided into the following three types according to their degree of virulence, which depends on HA activity: hyper-oral-toxic (HOT), intermediate, and non-HOT^8^. L-PTC/A^62A^, a non-HOT-type toxin, binds to glycoprotein-2 (GP2) on microfold (M) cells within the follicle-associated epithelium (FAE) of Peyer’s patches, and undergoes transcytosis across the epithelium^7^. By contrast, L-PTC/B^Okra^, a HOT-type toxin, is absorbed from the small intestine via enterocytes and M cells; however, the interaction between L-PTC/B^Okra^ and enterocytes remains poorly characterized. Here, we show that the M cell–independent entry route is pivotal for the higher oral toxicity of L-PTC/B^Okra^, and identify low-density lipoprotein (LDL) receptor–related protein 1 (LRP1) on enterocytes as a transcytosis receptor for L-PTC/B^Okra^.

## Results

### Uptake of L-PTC/B^Okra^ via enterocytes

To determine the entry route of the HOT-type toxin, fluorescently labeled L-PTC/B^Okra^ was administered into the mouse small intestine. The HA-containing toxin complex L-PTC/B^Okra^ (BoNT+NTNHA+HA) binds both enterocytes and M cells (Fig. 1a)^8^. Consistently, NAP/B^Okra^ (NTNHA+HA) associated with both cell types, whereas BoNT/B^Okra^ and M-PTC/B^Okra^ (BoNT+NTNHA) showed no detectable binding (Fig. 1b). Thus, L-PTC/B^Okra^ bound to the epithelium via HA. Although L-PTC/B^Okra^ is absorbed from the small intestine via enterocytes and M cells, it remains unclear which entry route is pivotal for its higher oral toxicity. To address this issue, we treated mice with an anti-RANKL monoclonal antibody to deplete their M cells (Fig. 1c)^7,9,10^. M-cell depletion slightly increased the resistance to L-PTC/B^Okra^ toxicity following oral administration (Fig. 1d). However, L-PTC/B^Okra^ remained at least 20-fold more toxic than L-PTC/A^62A^ in M cell–depleted mice (Fig. 1e). These findings suggest that the enterocyte-dependent route is pivotal for L-PTC/B^Okra^ entry into the host.

**Figure 1.**
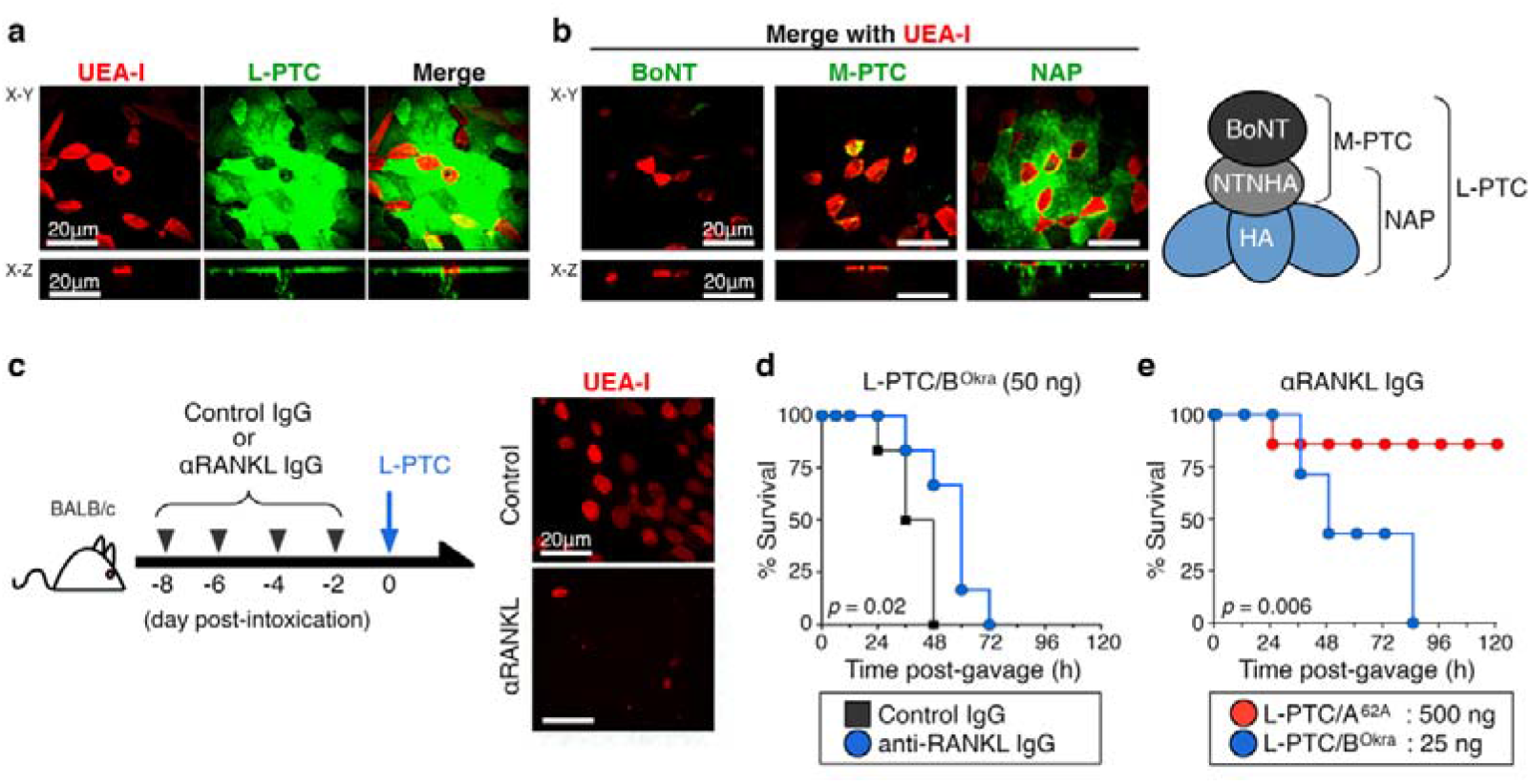
Host entry routes of the hyper-oral-toxic-type BoNT complex (L-PTC) from *C. botulinum* B-Okra in the intestinal epithelium. **a, b**, Representative images of whole-mount small intestine with AF488-labeled L-PTC (**a**) or other components (BoNT, M-PTC, NAP) (**b**) from B-Okra. M cells were visualized with rhodamine-labeled UEA-I (red). The upper and lower panels represent the X–Y and X–Z planes, respectively. Scale bar: 20 µm. **c**, A schematic model of M cell–depleted mice (left panel). BALB/c mice were treated with rat control IgG or rat anti-RANKL IgG (clone IK22-5) four times every 2 days. The M cells (red) were depleted within the FAE of Peyer’s patches in anti-RANKL IgG–treated mice (right panel). **d**, Survival curves of control or M cell–depleted mice (*n* = 6) that underwent intragastric administration of 50 ng of L-PTC/B^Okra^. **e**, M cell–depleted mice (*n* = 7) were challenged i.g. with L-PTCs from A-62A (500 ng, non-HOT) or B-Okra (25 ng, hyper-oral-toxic; HOT). **d, e**, Log-rank test.

### HA binds to LRP1 via *N*-glycans

HA facilitates BoNT attachment to the intestinal epithelium through its carbohydrate-binding activity^7,8^. HA is a hetero-dodecameric complex comprised of three subcomponents: HA1, HA2, and HA3 (also known as HA33, HA17, and HA70)^11,12^. HA1 and HA3 recognize galactose (Gal) and sialic acid (Sia) residues, respectively^12,13^ (Fig. 2a). We found that the HA1/B^Okra^ subcomponent had high affinity to villous epithelium and FAE, whereas HA2/B^Okra^ and HA3/B^Okra^ showed no detectable binding (Fig. 2a). These results show that HA1 is responsible for HA/B^Okra^ attachment to the intestinal epithelium.

**Figure 2.**
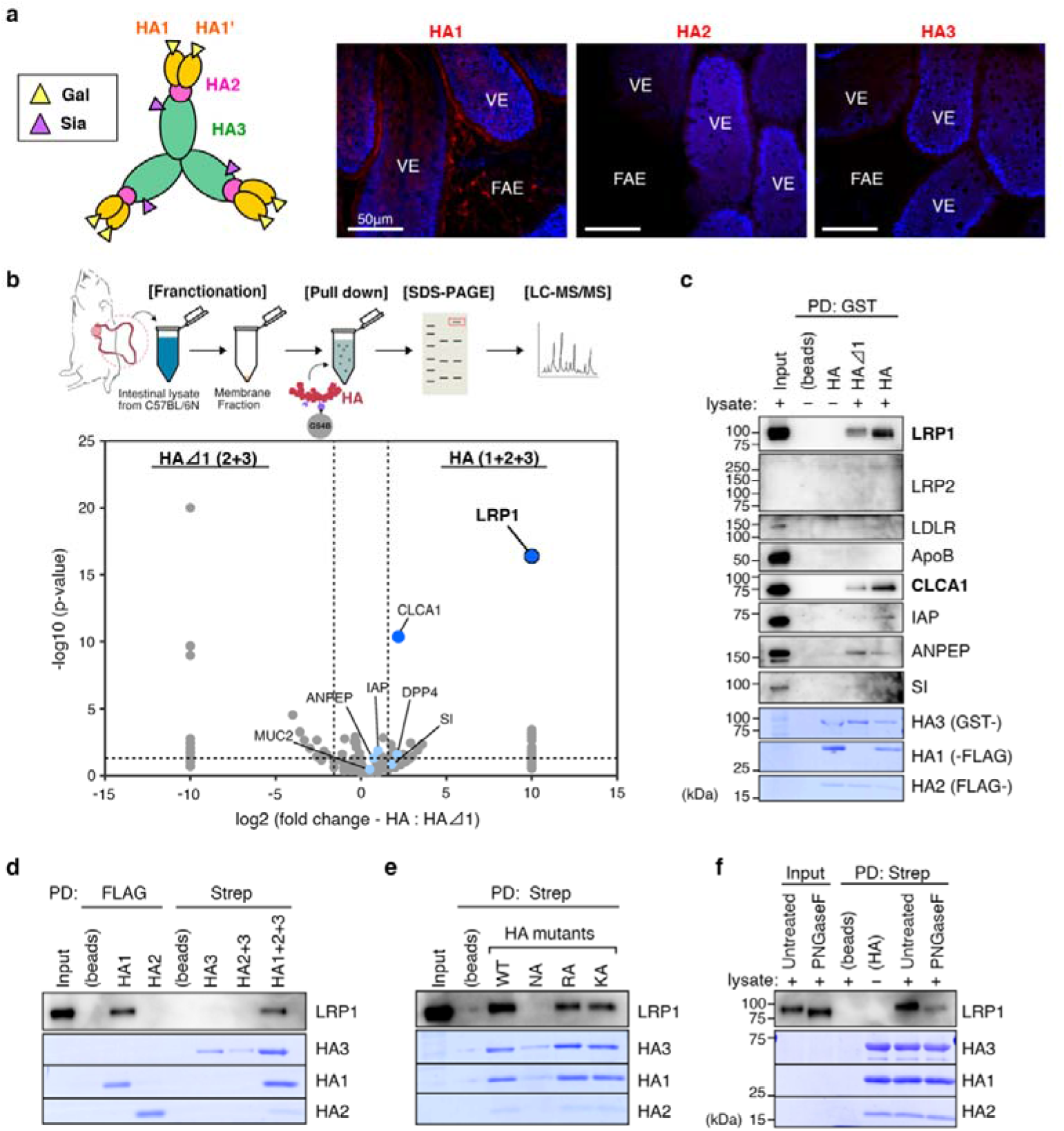
Identification of enterocyte transcytosis receptors for L-PTC/B^Okra^. **a**, A cartoon model of HA/B^Okra^ (left panel). The carbohydrate-binding sites are indicated by yellow (galactose, Gal) and purple (sialic acid, Sia) triangles. Representative images of whole-mount small intestine with AF568-labeled HA/B^Okra^ subcomponents (HA1, HA2, HA3) (right panel). VE, villous epithelium. Scale bar: 50 µm. **b**, Membrane fractions were extracted from IECs of mouse small intestine. The proteins interacting with HA1 were isolated from the lysates by GST pull-down with GST-HA/B^Okra^ (HA1+HA2+HA3), and identified by LC-MS/MS analysis. A volcano plot of −log_10_(*p*-value) versus −log_2_(fold change) depicts the 248 proteins identified. Negative control: GST-HAΔ1/B^Okra^ (HA2+HA3). **c**, GST pull-down assay with HA/B^Okra^ and the mouse intestinal lysates, followed by immunoblotting with antibodies against the identified proteins (LRP1, CLCA1, IAP, ANPEP, SI) and their associated proteins (LRP2, LDLR, ApoB). HA proteins were stained with Coomassie blue. **d**–**f**, Interaction between HA/B^Okra^ and LRP1 was analyzed by pull-down assay with the lysates prepared from Caco-2 cells. HA/B^Okra^ subcomponents (HA1-FLAG, FLAG-HA2, Strep-HA3, HA2+HA3, HA1+HA2+HA3) (**d**) and HA/B^Okra^ mutants (WT; NA, N286A in HA1; RA, R528A in HA3; KA, K607A in HA3) (**e**) were immobilized on FLAG M2 or Strep-Tactin beads. **f**, The cell lysates were treated with or without glycopeptidase F (PNGase F).

To identify entry receptors on enterocytes, we performed a glutathione S-transferase (GST) pull-down assay using HA/B^Okra^ (HA1+HA2+HA3) or HAΔ1/B^Okra^ (HA2+HA3) as bait, and the membrane fractions from mouse small intestine as prey (Fig. 2b and Fig. S1). Using HA, 41 proteins were enriched relative to HAΔ1 (Fig. 2b). Among these proteins, LRP1 and calcium-activated chloride channel regulator 1 (CLCA1) exhibited the highest HA/HAΔ1 fold change. Consistent with our mass spectrometry results, the pull-down assay showed that HA/B^Okra^ interacts with LRP1 and CLCA1 (Fig. 2c). LRP1 is a ubiquitously expressed membrane receptor protein (Fig. S2)^14^, whereas CLCA1 lacks a transmembrane segment and is abundantly secreted by goblet cells^15^. We therefore focused on LRP1 as a candidate transcytosis receptor for L-PTC/B^Okra^.

To clarify the HA–LRP1 interaction, we performed a pull-down assay using Caco-2 (human colon carcinoma) cell lysate as prey. Among the HA subcomponents, HA1, but not HA2 and HA3, interacted with human LRP1 (Fig. 2d). HA has three distinct binding sites that mediate direct interactions with Gal, Sia, and E-cadherin (Ecad)^12,13,16^; these interactions are abolished by the N286A (NA) substitution in HA1 and R528A (RA) and K607A (KA) substitutions in HA3, respectively^13^. Consistent with specific binding of HA1, HA/B^Okra^ wild type (WT), KA, and RA interacted with LRP1, whereas the NA mutant did not (Fig. 2e). Glycopeptidase F (PNGase F) treatment, which removes *N*-glycans from glycoproteins, diminished the binding of HA/B^Okra^ to LRP1 (Fig. 2f). Together, these results demonstrated that L-PTC/B^Okra^ binds to *N*-glycosylated LRP1 via HA1.

### Association of HA with LRP1-mediated lipoprotein transport

Using polarized monolayers of CMT-93 (mouse colon carcinoma) and Caco-2 cells as an *in vitro* transport model of intestinal epithelial cells, we found that HA/B^Okra^ attached to the apical cell surface and co-localized with multiple glycoproteins, including LRP1 (Fig. 3 and Fig. S3). Notably, HA/B^Okra^ exclusively co-localized with LRP1 in intracellular vesicles in both CMT-93 and Caco-2 cells (Fig. 3a and Fig. S3).

**Figure 3.**
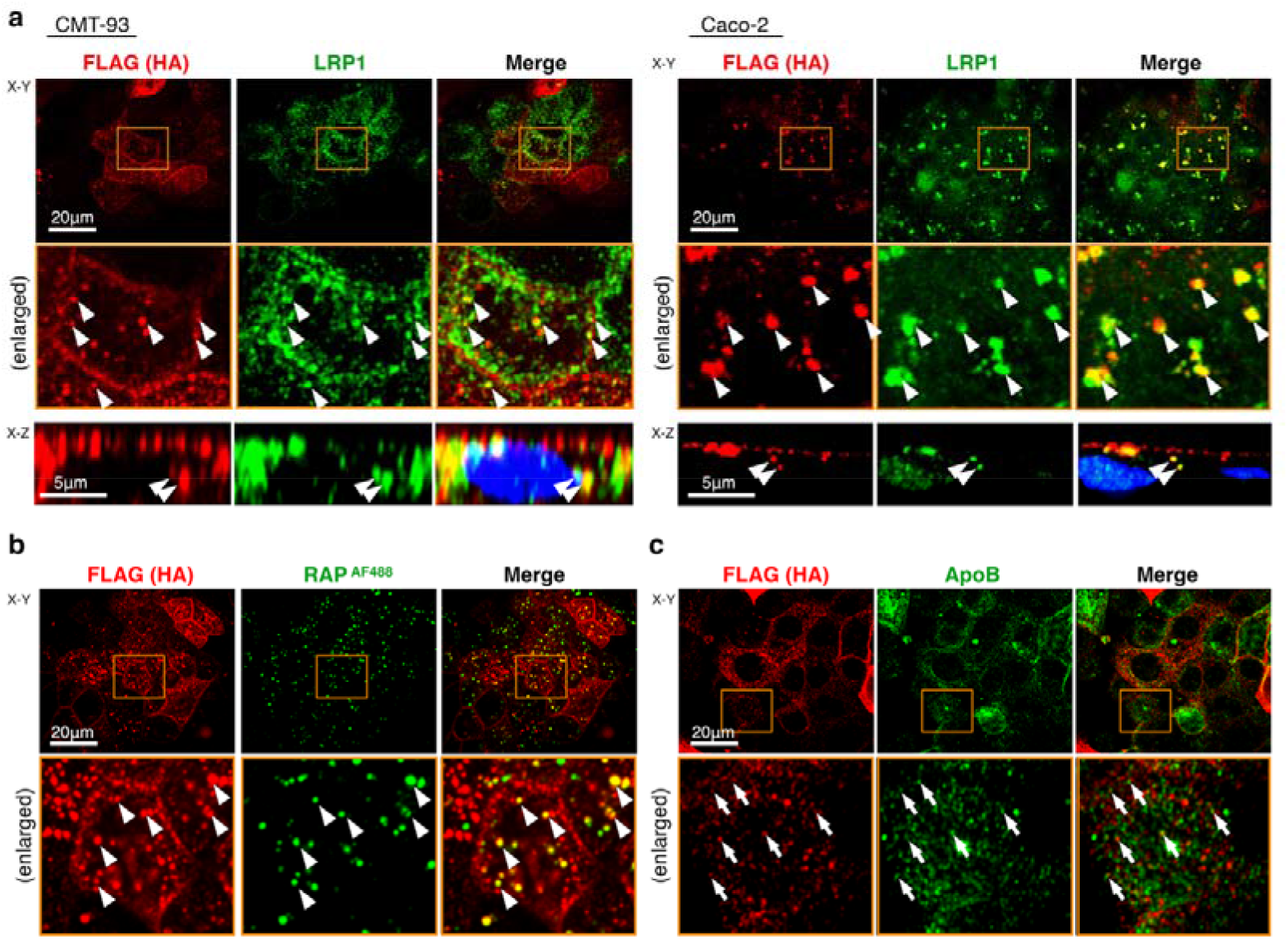
Co-localization of HA/B^Okra^ with LRP1 and associated proteins. **a**-**c**, Cell monolayers were apically treated with HA/B^Okra^ (20 nM, 2 h). **a**, Representative images of immunofluorescence staining with CMT-93 (left panel) and Caco-2 (right panel) cells. HA/B^Okra^ and LRP1 were probed by anti-FLAG (red) and anti-LRP1 (green) antibodies. **b, c**, CMT-93 cells were incubated with AF488-labeled GST-RAP (**b**) and probed with an anti-ApoB antibody (**c**). The lower panels of X–Y planes represent enlarged views of the corresponding boxed areas in the upper panels. The apical-basal axis is visualized in the X–Z plane. Arrowheads and arrows indicate co-localization and close proximity, respectively. Scale bar: 20 µm (X–Y), 5 µm (X–Z).

LRP1 is a member of the LDL receptor (LDLR) family of proteins and mediates the transport of chylomicrons containing triglycerides, phospholipids, cholesterol, and apolipoproteins^14^. Receptor-associated protein (RAP) serves as a chaperone that assists in the maturation of LRP-family proteins and can inhibit the binding of ligands such as lipoproteins^14^. However, recombinant GST-tagged RAP did not block HA/B^Okra^ uptake and co-localized with HA/B^Okra^ in intracellular vesicles (Fig. 3b and Fig. S4). HA-positive vesicles were accompanied by apolipoprotein B and E (ApoB and ApoE) and lipid droplets (Fig. 3c and Fig. S5), although HA did not directly interact with these components (Fig. 2b,c). These findings further support the model that the HA component of L-PTC/B^Okra^ enters enterocytes via LRP1.

### LRP1 acts as a transcytosis receptor for L-PTC/B^Okra^ in intestinal epithelial cells

Based on its expression pattern similarity to the LRP family *in vivo* (Fig. 2c, 4 and Fig. S6b, S7), we chose CMT-93 cells as a model to study the intestinal epithelium. To determine the functional impact of LRP1, we generated LRP1 knockout (KO) cells using a CRISPR/Cas9 system (Fig. 4a and Fig. S7). HA/B^Okra^ bound similarly to the apical surface of monolayers comprising three different CMT-93 cell types: WT, LRP1-KO, and a KO cell line expressing human LRP1 (LRP1-KO+hLRP1) (Fig. 4b). This binding was abolished by an NA mutation that results in defective Gal binding (Fig. 4b). We then used in-cell ELISA to assess the cellular uptake of HA/B^Okra^, which could be blocked by Dynasore, a dynamin-dependent endocytosis inhibitor (Fig. 4c). The uptake remained unaffected in LRP1-KO cells (Fig. 4c). Although LRP1 serves as apical surface receptor (Fig. S1c), the cell binding and subsequent endocytosis of HA/B^Okra^ were mediated not only by LRP1 but also by other glycoproteins. Finally, we performed apical-to-basal transcytosis and translocation assays using 0.4-µm-pore Transwell inserts. To evaluate transcytosis, we used an Ecad-binding–defective KA mutant because HA-WT compromises the integrity of epithelial cell layers by inhibiting Ecad function^13,16,17^. The deletion of LRP1 expression significantly reduced the transcytosis of HA/B^Okra^-KA, which was rescued by human LRP1 re-expression (Fig. 4d and Fig. S8a). Similarly, the translocation of L-PTC/B^Okra^-WT and HA/B^Okra^-WT was significantly reduced in LRP1-KO cells (Fig. 4e and Fig. S8b,c). These findings demonstrated that LRP1 mediates the transcytosis of L-PTC/B^Okra^ via HA across the intestinal epithelial barrier in an apical-to-basal direction *in vitro*.

**Figure 4.**
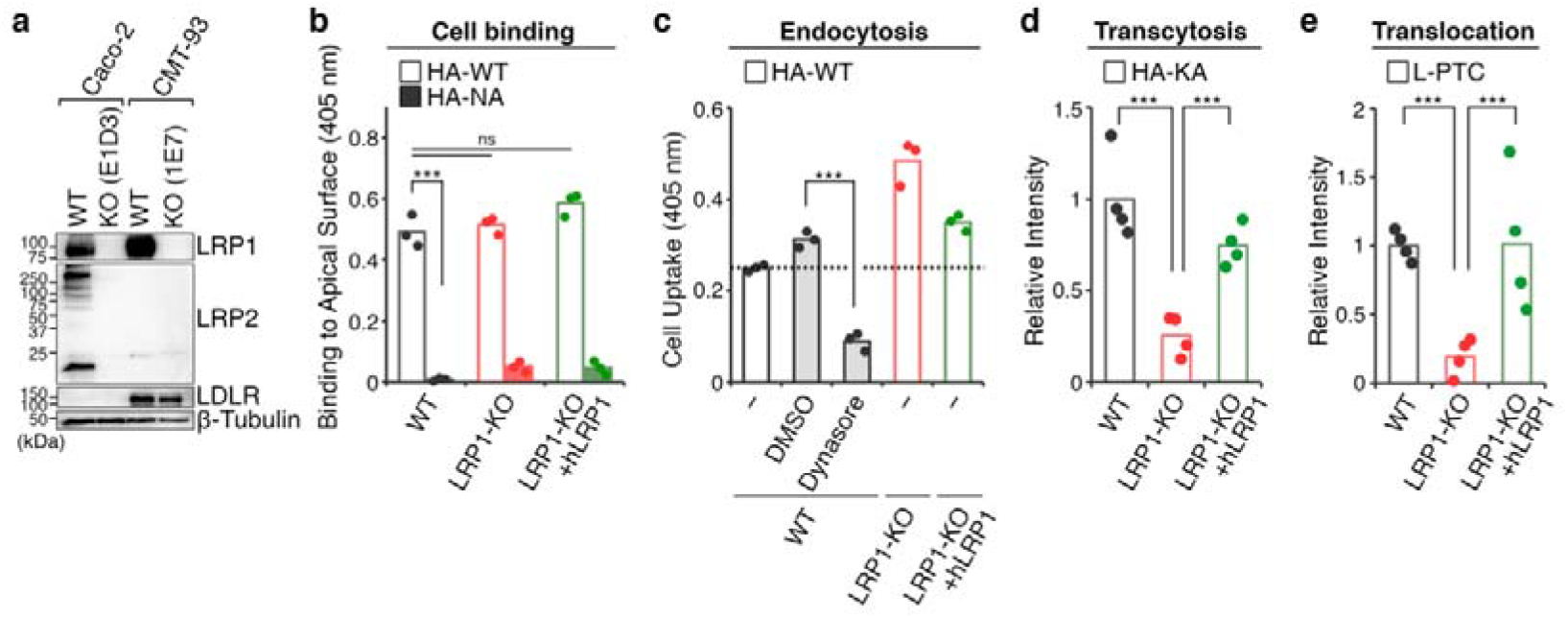
LRP1 is a transcytosis receptor for L-PTC/B^Okra^ in CMT-93 cell monolayers. **a**, Immunoblot analysis of the cellular expression of LRP-family proteins (LRP1, LRP2, LDLR) in Caco-2 (WT, LRP1/LRP2-KO) and CMT-93 (WT, LRP1-KO) cells. **b, c**, CMT-93 cell monolayers (WT, LRP1-KO, LRP1-KO+hLRP1) were apically incubated with HA/B^Okra^ at 4□ for 40 min (**b**) or 37□for 1 h (**c**). Binding (**b**) and endocytosis (**c**) of HA/B^Okra^ were analyzed by cell-ELISA and in cell-ELISA, respectively. **c**, Dynasore blocked the uptake of HA/B^Okra^. **d, e**, HA/B^Okra^-KA or L-PTC/B^Okra^-WT was added to the apical chamber of CMT-93 cells grown on Transwell membranes (0.4-µm pore diameter). After incubation at 37□for 24 h, proteins that had transcytosed or translocated into the basal media were collected by trichloroacetic acid precipitation, separated on SDS-PAGE, and detected by immunoblot analysis using anti-FLAG and anti-BoNT/B antibodies. Densitometry analysis was performed with ImageJ software to normalize FLAG-tagged HA1 and intact BoNT levels to the corresponding BSA levels. BSA was added to the media and used as an internal standard. Values represent the median of triplicate (**b, c**) or quadruplicate (**d, e**) wells. The data were analyzed by two-tailed student’s *t-*test (\*\*\**p* < 0.001, ns: not significant).

### LRP1 mediates BoNT uptake via enterocytes

We found that LRP1 is highly expressed in enterocytes within villous epithelium and resides primarily around nuclei, with some apical and basal distribution (Fig. 5a). AF568-labeled HA/B^Okra^ bound to the luminal surface of enterocytes and was internalized into LRP1-containing vesicles (Fig. 5a). These vesicles were accompanied by ApoB and ApoE (Fig. 5b and Fig. S9). These results confirmed that the HA of L-PTC/B^Okra^ acts via the LRP1-mediated lipoprotein transport system in mouse small intestine.

**Figure 5.**
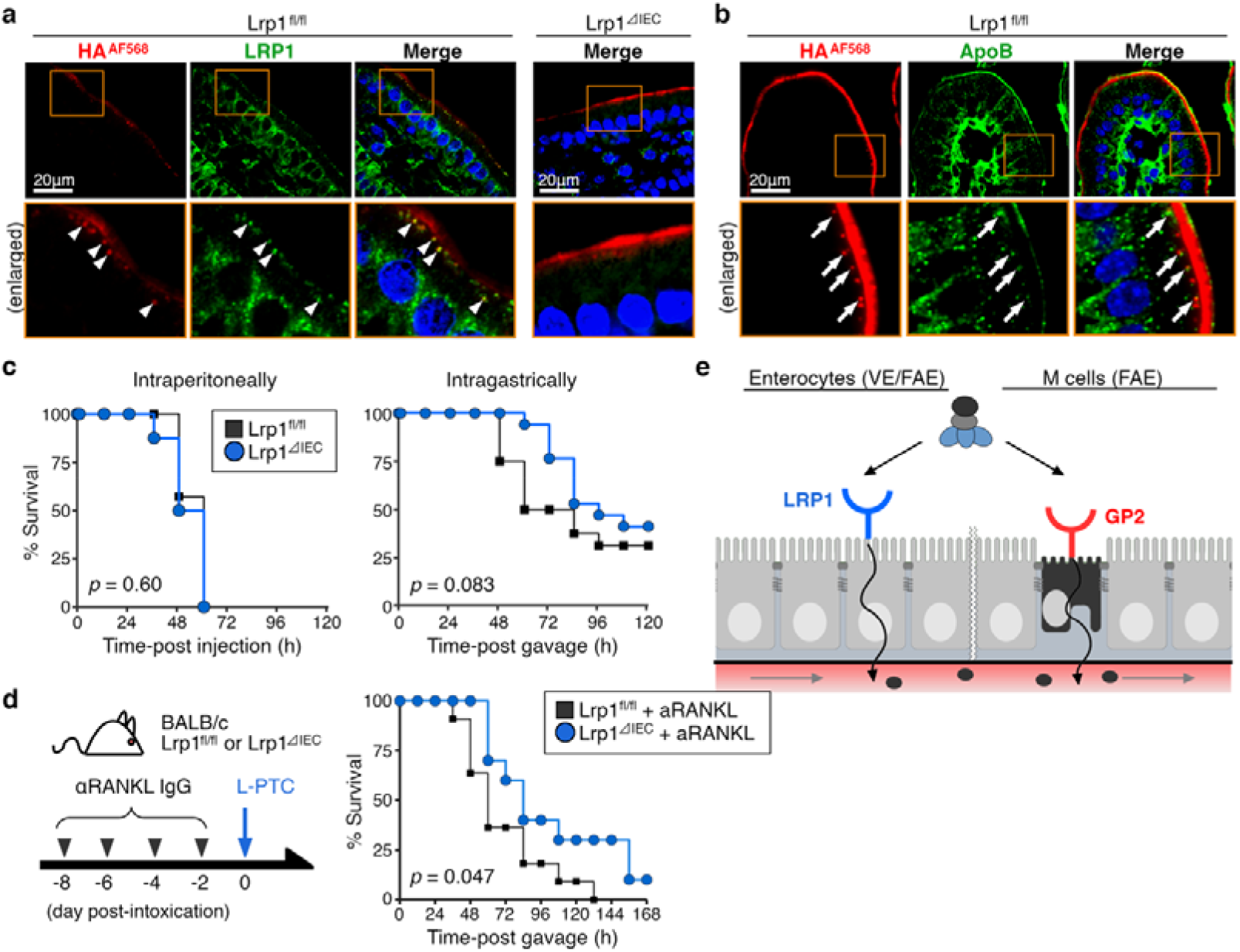
LRP1 facilitates intestinal absorption of BoNT/B^Okra^ in mice. **a, b**, Representative images of fixed frozen duodenal sections with AF568-labeled HA/B^Okra^ (red) from control (*Lrp1*^fl/fl^) and intestinal epithelial cell–specific LRP1-deficient (*Lrp1*^ΔIEC^; *Lrp1*^fl/fl^ Vil-Cre) mice. Nuclei, LRP1, and ApoB were visualized with Hoechst 33342 (blue), anti-LRP1 (green), and anti-ApoB antibodies (green), respectively. The lower panels represent enlarged views of the corresponding boxed areas in the upper panels. Arrowheads and arrows indicate co-localization and close proximity, respectively. Scale bar: 20 µm. **c**, *Lrp1*^fl/fl^ and *Lrp1*^ΔIEC^ mice were challenged with L-PTC/B^Okra^ i.p. (left panel, 25 pg, *n* = 7–8) and i.g. (right panel, 10 ng, *n* = 16–17). **d**, A schematic model of M cell–depleted mice (left panel). *Lrp1*^fl/fl^ and *Lrp1*^ΔIEC^ mice were treated with rat anti RANKL IgG (clone IK22-5) four times every 2 days. Survival curves of the M cell-depleted mice after intragastric administration with 15 ng of L-PTC/B^Okra^ (right panel, *n* = 10–11). **e**, A schematic model of BoNT absorption from the intestine via enterocytes and M cells. **c**, Gehan–Wilcoxon test. **d**, Log-rank test.

Given that germline KO of LRP1 is embryonic lethal in mice due to defects in embryo^18^, we generated intestinal epithelial cell (IEC)-specific *Lrp1*-deficient (*Lrp1*^ΔIEC^) mice by crossing *Lrp1*-floxed (*Lrp1*^fl/fl^) mice with Villin-Cre (Vil-Cre) transgenic mice (Fig. S6a). LRP1 expression in enterocytes was abolished in *Lrp1*^ΔIEC^ mice (Fig. 5a and Fig. S6b). *Lrp1*^fl/fl^ and *Lrp1*^ΔIEC^ mice exhibited comparable susceptibility to intraperitoneal injection of L-PTC/B^Okra^ (Fig. 5c). By contrast, *Lrp1*^ΔIEC^ mice showed a trend toward increased resistance to L-PTC/B^Okra^ toxicity compared with *Lrp1*^fl/fl^ mice (Fig. 5c). Upon M-cell depletion by administration of an anti-RANKL antibody, *Lrp1*^ΔIEC^ mice showed greater resistance to L-PTC/B^Okra^ toxicity than *Lrp1*^fl/fl^ mice (Fig. 5d). Taken together, these findings indicate that LRP1 facilitates BoNT absorption via enterocytes, in addition to GP2-dependent uptake via M cells, in the small intestine (Fig. 5e and Fig. S10).

## Discussion

The toxicities of all orally administered BoNT serotypes are augmented by NAPs^2^ and OrfX proteins^19^. Notably, HA facilitates BoNT absorption from the epithelium in the small intestine^2,7,8^, and we previously proposed that the toxicity of orally administered L-PTC can be determined by the type of HA rather than the BoNT serotype^8^. L-PTC from *C. botulinum* A-62A (L-PTC/A^62A^, non-hyper-oral-toxic type^2,8^) enters the host by exploiting intestinal M cells^7^. On the other hand, L-PTC from B-Okra (L-PTC/B^Okra^, hyper-oral-toxic type^2,8^) can enter through both M cells and enterocytes^8^. Enterocytes are the predominant epithelial cell type in the small intestine, whereas M cells are restricted to the FAE of Peyer’s patches and account for 5–10% of FAE cells^20^. Consistent with this cellular distribution, our results indicate that the M cell route is associated with, but not responsible for the higher oral toxicity; instead, entry through enterocytes is the major route of the toxin uptake.

In M cells, GP2, an uptake receptor for a subset of commensal and pathogenic bacteria^21,22^, serves as a transcytosis receptor for both L-PTC/A^62A7^ and L-PTC/B^Okra^ (Fig. S10). However, GP2 is exclusively expressed on M cells, indicating that an additional transcytosis receptor is required for toxin entry through enterocytes. In this study, we identified LRP1 as an enterocyte entry receptor for L-PTC/B^Okra^.

LRP1 (also known as CD91 or α2-macroglobulin receptor) is a glycosylated single-transmembrane protein belonging to the LRP family^14^. The pro-protein (600 kDa) of LRP1 is proteolytically nicked by furin, resulting in a 515 kDa α-chain and an 85 kDa β-chain (Fig. S2). The α-chain has four clusters (clusters I–IV) of ligand-binding repeats that interact with more than 70 distinct ligands^14,23–25^. Notably, LRP1 has been reported as a potential receptor for various pathogens, including bacterial toxins such as *Clostridium perfringens* TpeL^26^, *Pseudomonas* exotoxin A^27^, *Clostridium novyi* alpha toxin^28^, *Clostridioides difficile* toxin A^29^, *Helicobacter pylori* vacuolating cytotoxin^30^, and *Pasteurella multocida* toxin^31^, along with plant toxins such as ricin^32^ and viruses such as Rift Valley fever virus^33^. These interactions are blocked by RAP, a molecular chaperone of LRP1^26,27,31,32^. By contrast, the addition of GST-RAP did not hinder the interaction between HA/B^Okra^ and LRP1 (Fig. 3b and Fig. S4, S5). Unlike other ligands involved in protein–protein interactions, HA/B^Okra^ binds to LRP1 via *N*-glycans (Fig. 2f). LRP1 has 39 potential *N*-glycosylation sites in the extracellular domain (Fig. S2). Aminopeptidase N (ANPEP), sucrase–isomaltase (SI), intestinal alkaline phosphatase (IAP), and dipeptidyl peptidase 4 (DPP4) are membrane-bound glycoproteins and brush border membrane markers^34^. However, these glycoproteins had a lower affinity to HA/B^Okra^ (Fig. 2b, c) and failed to mediate HA/B^Okra^ endocytosis (Fig. S3). We believe that the triskelion-shaped HA (470 kDa) armed with six HA1 (Fig. 2a)^11,12^ effectively recognizes the highly *N*-glycosylated LRP1 (α-chain: 515 kDa) through a multivalency effect^35^. The carbohydrate-binding activity of HA is robust under acidic (pH 6.0) and near-neutral (pH 7.4) conditions^8,36^. Thus, HA can interact with LRP1 in endocytic vesicles despite rapid acidification, which may allow the toxin complex to remain associated with the receptor during transcytosis.

Dietary lipids are absorbed from small intestinal enterocytes and incorporated into nascent chylomicrons (CMs)^37^. CMs are then secreted into lymph and subsequently hydrolyzed in the circulation, allowing the delivery of free fatty acids to peripheral tissues. LRP-family proteins mediate the secretion and transport of lipoproteins. In enterocytes, LRP1 binds to ApoB-48 and ApoE of CMs^24,25^ and mediates exocytosis^38^. HA/B^Okra^ binds to LRP1 in a manner distinct from lipoproteins^39,40^, and in this study, it was accompanied by ApoB, ApoE, and lipid droplets and co-localized with LRP1. These findings suggest that L-PTC/B^Okra^ may exploit the lipoprotein transport system for transcytosis to the basal lamina.

Taken together, our findings demonstrate that LRP1 is a major enterocyte transcytosis receptor for L-PTC/B^Okra^. This study elucidated the mechanism by which hyper-oral-toxic (HOT)-type BoNT crosses the intestinal epithelium to enter the host. Following transcytosis to the basal side, HA of L-PTC/A^62A^-WT and L-PTC/B^Okra^-WT further augments oral toxicity by disrupting the E-cadherin– mediated epithelial barrier^7,16,17,41,42^. Understanding this molecular mechanism will serve as a foundation for the development of more effective therapeutic strategies and novel drug delivery systems that facilitate drug transport across the epithelial barrier in the small intestine.

## Acknowledgments

We thank Chiyoko Aoki, Yuki Sano, Yuriko Tanaka, Sachiyo Akagi, Hitomi Kuraoka, Yuki Konoshita, and Hiromi Honda for providing technical assistance, and the members of the Fujinaga laboratory for valuable discussions. We thank Kazunobu Saito (Osaka University, RIMD) for mass spectrometric analysis, and Tetsuichiro Inai (Fukuoka Dental College) for the gift of CMT93-I cells. We thank Editage (www.editage.com) for English language editing. S.A. was supported by JSPS KAKENHI Grant Numbers 15J03770, 19K21257, 20K16240, and 23K14517. T.M. was supported by JSPS KAKENHI Grant Numbers 16K19123, 18K07107, and 25K10359. Y.F. was supported by JSPS KAKENHI Grant Numbers 24K02278.

## Author contributions

S.A., T.M., and Y.F. conceived the project idea and designed the experiments. S.A. and T.M. executed the experiments and performed the data analysis. C.M. assisted in the CRISPR/Cas9 knockout experiments. H.Y. provided the anti-RANKL antibody. K-a.I. and T.T. provided *Lrp1*-floxed mice. T.K. and H.O. provided Vil-Cre mice. N.K. and K.H. assisted with the whole-mount immunostaining experiments. M.Z. assisted with data curation and resource. S.A. and Y.F. wrote the manuscript with editorial input from all of the authors.

## Data availability

All experimental data are contained within the article.

### Additional Information Supplementary information

This article includes:

Figs. S1 to S11

